# Cytokine gene polymorphism and parasite susceptibility in free-living rodents: importance of non-coding variants

**DOI:** 10.1101/2021.09.16.460687

**Authors:** Agnieszka Kloch, Ewa J Mierzejewska, Renata Welc-Falęciak, Anna Bajer, Aleksandra Biedrzycka

## Abstract

Associations between genetic variants and susceptibility to infections have long been studied in free-living hosts to infer contemporary evolutionary forces shaping genetic polymorphisms of the immunity genes. Despite extensive studies of receptors, such as MHC or TLR, little is known about efferent arm of the immune system. Cytokines are signalling molecules that trigger and modulate the immune response, acting as a crucial link between innate and adaptive immunity. In the present study we investigated how genetic variation in cytokines affects susceptibility to parasitic diseases in bank voles. We focused on three cytokines: tumour necrosis factor (TNF), lymphotoxin alpha (LTα), and interferon beta (IFNβ1). Two SNPs in LTα and two in IFNβ1 significantly affected susceptibility to nematodes, and was of them was also associated with susceptibility to microbial pathogen *Bartonella*. All these variants displayed signatures of selection. One of the variants was synonymous, and one was located in an intron. Our study shows that cytokines are prone to parasite-driven selection, and non-coding variants may play an important role in susceptibility to infections in wild systems.

## 1. Introduction

Parasite-driven selection is considered a key factor shaping the evolution of the components of the immune system. In mammals and higher vertebrates, the immune system comprises dozens of interacting molecules, yet the mechanisms of this selection have been comprehensively studied only in the case of major histocompatibility complex (MHC) genes (e.g. Radwan et al 2020). Though the MHC plays an important role in the antigen-based pathogen recognition, it is not the only nor the main factor responsible for resistance against pathogens (Jepson et al. 1997). Recently, researchers focused on the components of the innate immunity, in particular the toll-like receptors (eg. Fornuskova et al. 2013, Babik et al. 2015, Kloch et al. 2018) but evolution of other elements of the immune system, including cytokines, remains poorly understood.

The cytokines, signalling molecules capable of triggering and modulating the immune response are the crucial link between innate and adaptive immunity. As an efferent arm of the immune system and they are expected to be more evolutionary constrained than elements of the afferent arm (Chapman et al. 2016) but a handful of studies reported signatures of balancing or positive selection in this group of molecules. For instance, balancing selection was found in interleukins Il-1B, Il-2, and TNF in field voles (Turner et al. 2012), and in Il-10 and CD14 in humans (Ferrer-Atmetlla et al. 2008). Another evidence for contemporary parasite-driven selection operating on cytokines are significant associations between genetic variants and susceptibility to infections. In humans, polymorphism within the LTα has been linked to several diseases, such as *Mycobacterium leprae* (Ware 2005) and malaria (Barbier et al. 2008). Among rodents, both positive and negative associations between variation in interleukins and infections with uni- and multicellular parasites were find in the field vole (Turner et al. 2011). Single study from a free-living animal revealed that variants affecting susceptibility may be located outside the coding part of a cytokine gene (Guivier et al. 2010) but further studies are needed to evaluate the role of non-coding polymorphisms.

To better understand the role of the parasite-driven selection in maintaining polymorphism in cytokines, and to further explain the role of this polymorphism in resistance against pathogens in the wild, we studied three cytokine genes: tumor necrosis factor (TNF), lymphotoxin alpha (LTα) formerly known as tumor necrosis factor beta (TNFβ), and interferon beta (IFNβ1). Their versatile role within the immune system, along with an evidence from human studies underlying its importance in a response against various pathogens, makes them a promising candidate gene for studying the mechanisms of parasite-driven selection. TNF initiates acute phase response and acts as an endogenous pyrogen contributing to the inflammation. Through stimulating endothelial cells in blood vessels it plays a role in preventing the pathogen from entering the bloodstream and containment of local infection (Murphy et al. 2012). LTα is expressed in lymphocytes and it plays a major role in immunomodulation and signal transduction within the immune system. It also induces inflammation and is the key factor facilitating the innate immune response through activation of IFNβ and NF-κB pathways (Iizuka et al. 1999; Ware 2005). LTα is also necessary for effective adaptive responses involving T and B lymphocytes (De Togni et al. 1994). IFNβ is produced in response to viral but also bacterial infections (Nagarajan 2011). Through specific pathway, it plays a major role in linking innate and adaptive immunity. We expected to detect signatures of selection acting on the studied loci and to find significant associations between specific variants and susceptibility to infections.

## 2. Material and methods

### 2.1. Samples and parasite screening

In the present work we used samples collected in 2005 and in 2016 in three sites in NE Poland (Urwitałt, Pilchy, Tałty) located within 30 km distance. There are temporal and spatial differences in parasite prevalence between these sites described comprehensively elsewhere (eg. Bajer et al. 2005, Welc-Faleciak et al. 2008), and associations between parasite load and variation in MHC and TLR genes have been reported (Kloch et al. 2010, 2018). Number of samples per year per site is given in Table S1.

The field procedures followed the guidances of the National Ethics Committee for Experimentation on Animals and were approved by the Local Ethical Committee no. 1 in Warsaw, decisions no. 280/2003 and 304/2012. Field procedures and analysis of helminth infections followed protocols described in Bajer et al. (2005) and Kloch et al. (2010). In brief, voles were live-trapped in wooden traps, transported to the field station, euthanised, and sectioned in search for internal helminths. Infections with the intestinal protist *Cryptosporidium* sp. were identified in faecal smears using the Ziehl-Nielsen staining technique (Heinriksen and Pohlenz 1981). Blood pathogen *Bartonella sp*. was identified by PCR using primers described by Norman et al. (1995), and for *Babesia microti* we used protocol described by Persing et al. (1992). Details of the PCR reactions are given in Table S2.

### 2.2. Sequencing and genotyping

To minimize PCR errors, cytokine genes were amplified using high fidelity polymerase Phusion or Q5 (NewEngland Biolabs). The mix contained 10uM dTNPs, 0.5uM of each primer, 0.02U/ul of the polymerase, and 10-50 ng of genomic DNA. Primer sequences and PCR conditions specific for each locus are given in Table S2. Amplified regions spanned most of the expressed exons and respective separating introns (Table S3).

Samples from 2005 and 2016 were processed separately following the same protocol. In both experiments, the amplicons were pooled for each individual as described previously (Kloch et al. 2018) and purified twice using CleanUp kit (Aabiot). The libraries were constructed using using Nextera XT DNA Library Preparation Kit, and sequenced using MiSeq Reagent Kit v3 in Illumina MiSeq machine. The only difference between experiments was the number of cycles used for sequencing: samples from 2005 were sequenced using 150 cycles, and samples from 2016 using 300 cycles. Both runs gave high coverage exceeding 1000x per site and per sample.

Reads from both runs were processed in the same pipeline as described previously (Kloch et al. 2018). In brief, adapter-clipped reads were mapped using bwa-mem ver. 0.7.12 with default parameters (Li 2013) against reference constructed from regions including the genes of interests extracted from bank vole genome (PRJNA290429). Duplicated read-pairs were removed using Picard MarkDuplicates (http://picard.sourceforge.net), and variants were called in two-step procedure in FreeBayes v 1.1.0-60 (Garrison & Marth 2012). In the first round, potential variants were called using the following parameters: minimal fraction of alternate allele of 20% as recommended by Nielsen et al. (2011), minimum number of reads supporting alternate allele > 2, and minimal read coverage > 5. The results were filtered using vcffilter v. 41 (https://github.com/ekg/vcflib) with following conservative criteria: QUAL/AO > 10 & DP > 10 & SAF > 0 & SAR > 0 & RPR > 0 & RPL > 0 resulting in high confidence variants. They were used to construct haplotypes in a second round of variant calling in FreeBayes were physical position on a read was used for phasing by specifying --max-complex-gap 37. The remaining SNPs were phased computationally using PHASE (Stephens et al. 2001), and phased alleles were reconstructed using vcfx (Castelli et al. 2015, Lima et al. 2016). The reading frames and intron/exon structure were resolved through an alignment to the orthologue mouse sequences. The sequencing revealed 43 SNP in ~2000bp in total.

### 2.3. Associations between genetic variants and parasite load

The sample size was strongly limited by the permission from the Ethical Committee. Thus, in order to minimize false positives we selected such a combination of variables fitted in the models that produce reliable results with lowest possible sample size (Hong and Park 2012).

First, we removed variants and individuals with missing SNPs. Next, SNPs were filtered in PLINK v1.90 (Purcell et al. 2007) so that only those in Hardy-Weinberg equilibrium (threshold of p<0.001) and in linkage equilibrium (r>0.7) were retained. (We tested for LD values from r>0.6 to r>0.9 and all the values resulted in the same set of SNPs). This ensured that genetic variants fitted in models can be treated as independent explanatory variables. After filtering, of the initial 18 SNPs in TNF, 17 in LTα, and 8 in IFNβ1 we retained one SNP in TNF, 7 SNP in LTα, and two in IFNβ1.

In a second step, we excluded from the models variants with MAF<0.5 and those present in fewer than 5 animals (see Table S4), as we lacked statistical power to test for their effects. As response variables we used only pathogens that infected 20-80% of hosts (Table S5).

We started the modelling from testing for the effect of non-genetic terms that may affect the parasite load: year and site of sampling, and host sex and host age approximated by its body mass (Table S6). If p<0.1, these terms were included in the models as co-factors. For each studied gene, we constructed two sets of generalized linear models (GLM) using R with either i) parasite presence/absence, or ii) parasite abundance (number of parasites of a given species per host) as the response variables. We used binomial errors for parasite presence/absence and Poisson for abundance, controlling for overdispersion. The explanatory variables were SNPs retained after filtering and non-genetic terms as described above. To minimize type I error when comparing effects of more than one genetic term, the p-values were corrected for multiple comparisons using Benjamini-Yekutieli false discovery rate (Benjamini and Yekulteli 2005) using p.adjust function in R. For all the calculations we used R version 3.6.3 (R Core Team 2020).

### 2.4. Tests of selection

To identify potential targets of selection, we computed phylogenetically controlled codon-based tests which are suitable for identifying sites under selection using sets of sequences from a single species (Kosakovsky Pond and Frost 2005). For the calculations, we used DataMonkey server (Delport et al. 2010). Prior to analysis, we tested for possible recombinations using GARD, Genetic Algorithm Recombination Detection (Kosakovsky Pond et al. 2006). We computed three models: MEME (Mixed Effects Model of Evolution) aimed at detecting episodic positive selection (Murrell et al. 2012); FUBAR (Fast Unconstrained Bayesian Approximation) detecting sites under pervasive diversifying selection (Murrell et al. 2013); and FEL (Fixed Effects Likelihood) that assumes the constant selection pressure for each site and also detects pervasive diversifying selection (Kosakovsky Pond and Frost 2005).

## 3. Results

### 3.1. Cytokine polymorphisms and susceptibility to infections

The only SNP in TNF gene retained after filtering did not affect presence/absence nor intensity of infection with any of the studied parasites (Table S7).

Two SNPs in LTα were significantly associated with parasite load (Table 1). Individuals with genotype CT in position 322 coding for non-synonymous substitution Gln > Arg were more often infected with the nematode *A. tianjinensis* than homozygotes TT (57.1 vs. 34.4% infected, respectively, Figure 1, Table 1). The alternative homozygote CC was found in a single host so we lacked statistical power to estimate its effect. An intronic variant LTα 525 affected abundance of infection with the nematode *H. mixtum*. Homozygotes TT were infected by fewer worms compared to genotypes GG and TG (Figure 1).

**Figure 1.**
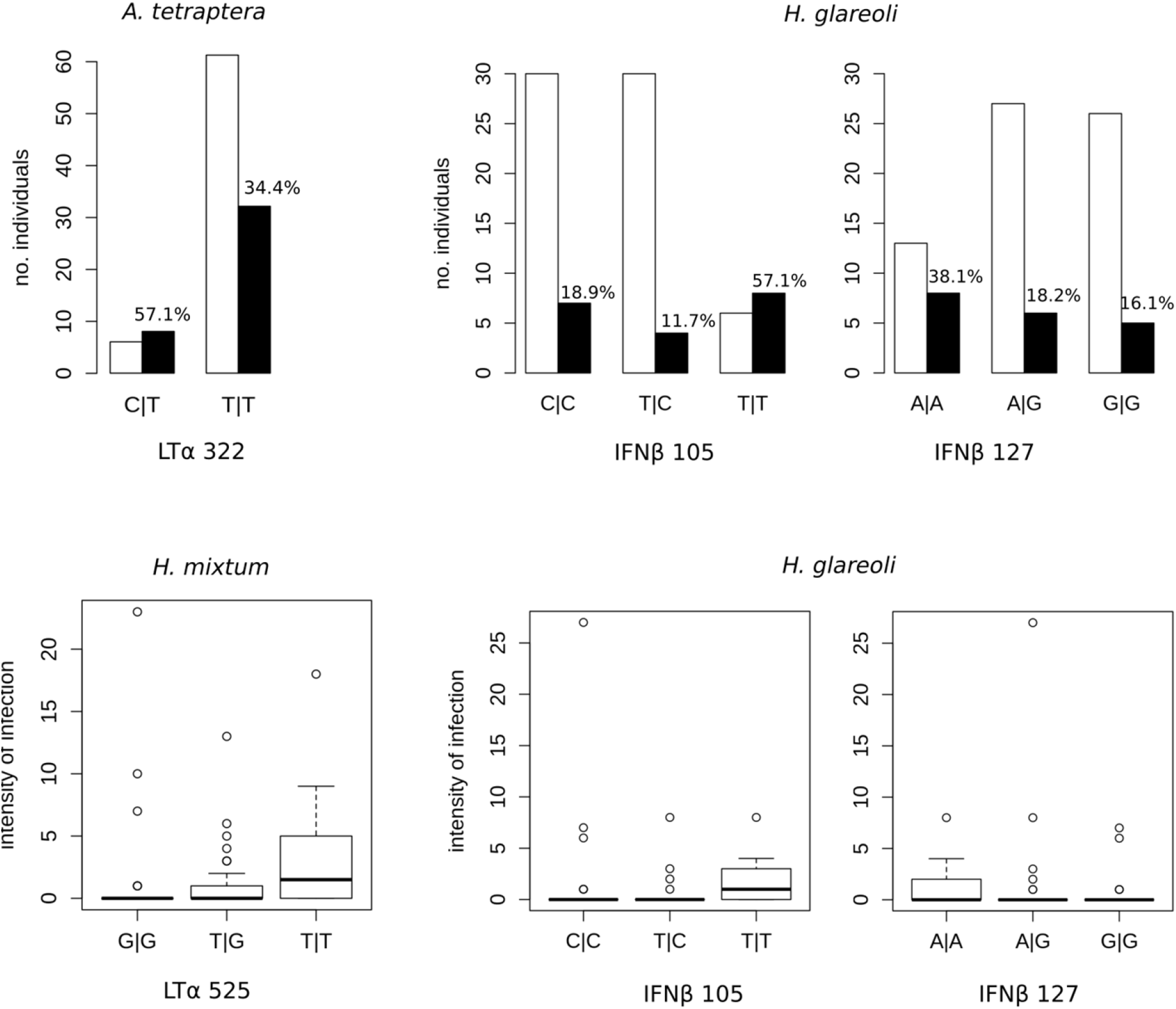
Effect of SNP genotype in a) LTα and b) IFNβ on the risk of infection and parasite abundance. % of infected voles is given in graphs where the allele affected the risk of infection.

**Table 1.**
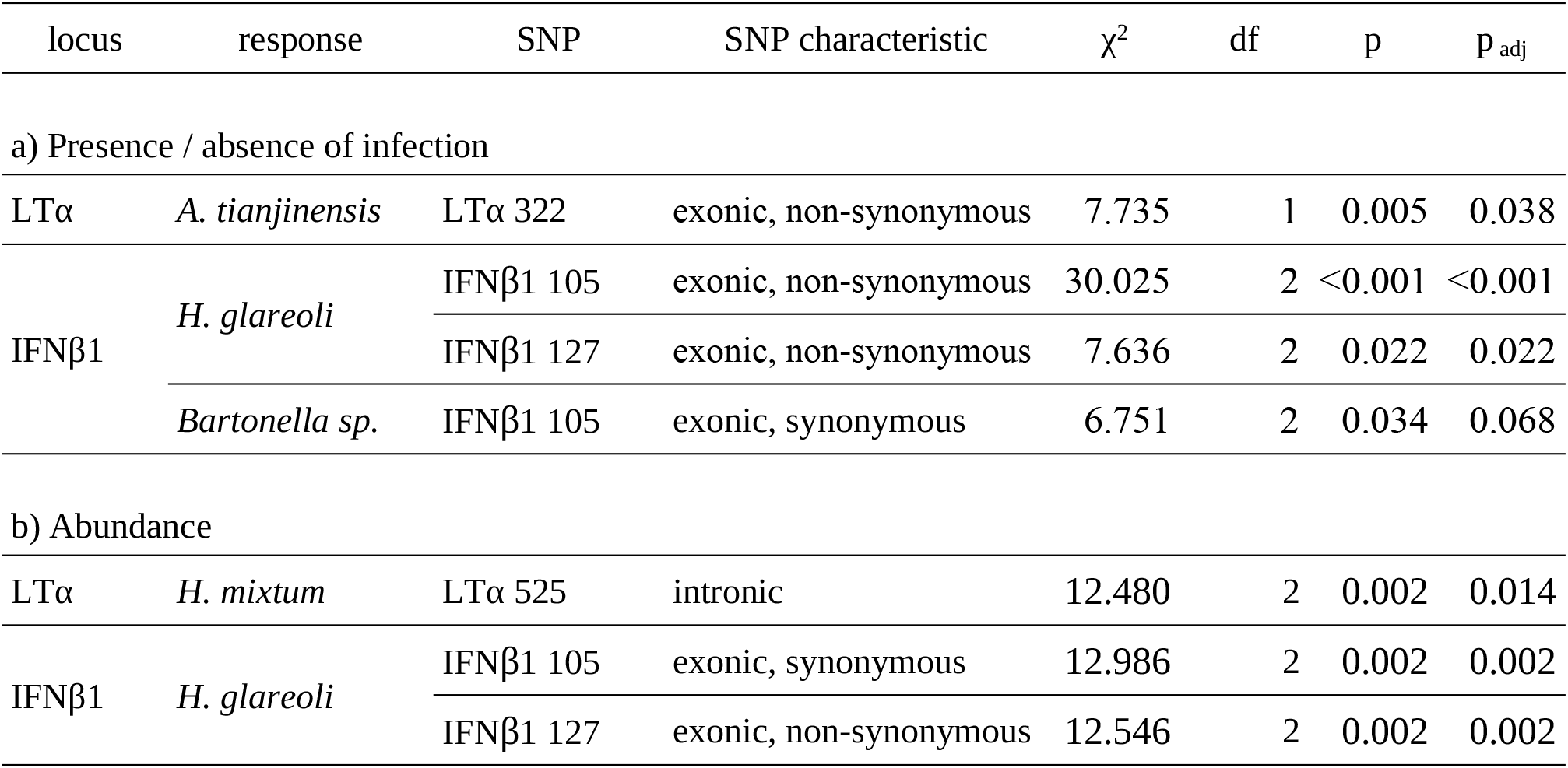
Summary of GLM models showing significant effect of SNP genotype at given locus on the parasite load. Full models are given in Table S7. p_adj_ denotes p-value adjusted for multiple comparisons.

Two IFNβ1 variants affected the parasite load (Table 1). Voles heterozygous in synonymous variant IFNβ1 105 were less frequently infected with the nematode *H. glareoli* (11.7%) compared to homozygotes TT (57.14%) and CC (18.9%, Figure 1), and they also harboured fewer worms (0.411) compared to 1.78 vs 1.19 in the respective homozygotes. In IFNβ1 127 homozygotes AA were most often infected (38%), and prevalence was similar in genotypes AG and GG. Homozygotes GG had the lowerst infection intensity but average high parasite load in heterozygotes may be contributed to a single highly infected individual. When it was excluded, intensity of infection was similar in genotypes AG and GG.

Genotype in IFNβ1 105 affected also infection risk with blood parasite *Bartonella* yet this effect was marginally non-signficant after controlling for multiple comparisons (p=0.068, Table 1). Heterozygous voles were more often infected than either homozygote (36.3% compared to 18.4% in CC and 21.24% in TT.

### 3.2. Signatures of selection

Selection tests were generally consistent across loci, and each of them reported the same set of codons (Table S8). FUBAR and FEL indicated 8 codons under pervasive positive selection in TNF, 7 in LTα, and one in IFNβ1 (Figure 2). Among these codons, four in TNF and three in LTα displayed signatures of episodic positive selection, as indicated by MEME model. Positively selected codon 525 in LTα was associated with increased prevalence of *H. mixtum*, and positively selected codon compring variant IFNβ1 127 was associated with higher prevalence and intensity of *H. glareoli*. Two codons in TNF and one in IFNβ1 displayed signatures of negative selection. The negatively selected codon comprising variant IFNβ1 105 was associated both with higher prevalence and intensity of *H. glareoli* and lower prevalence of *Bartonella*.

**Figure 2.**
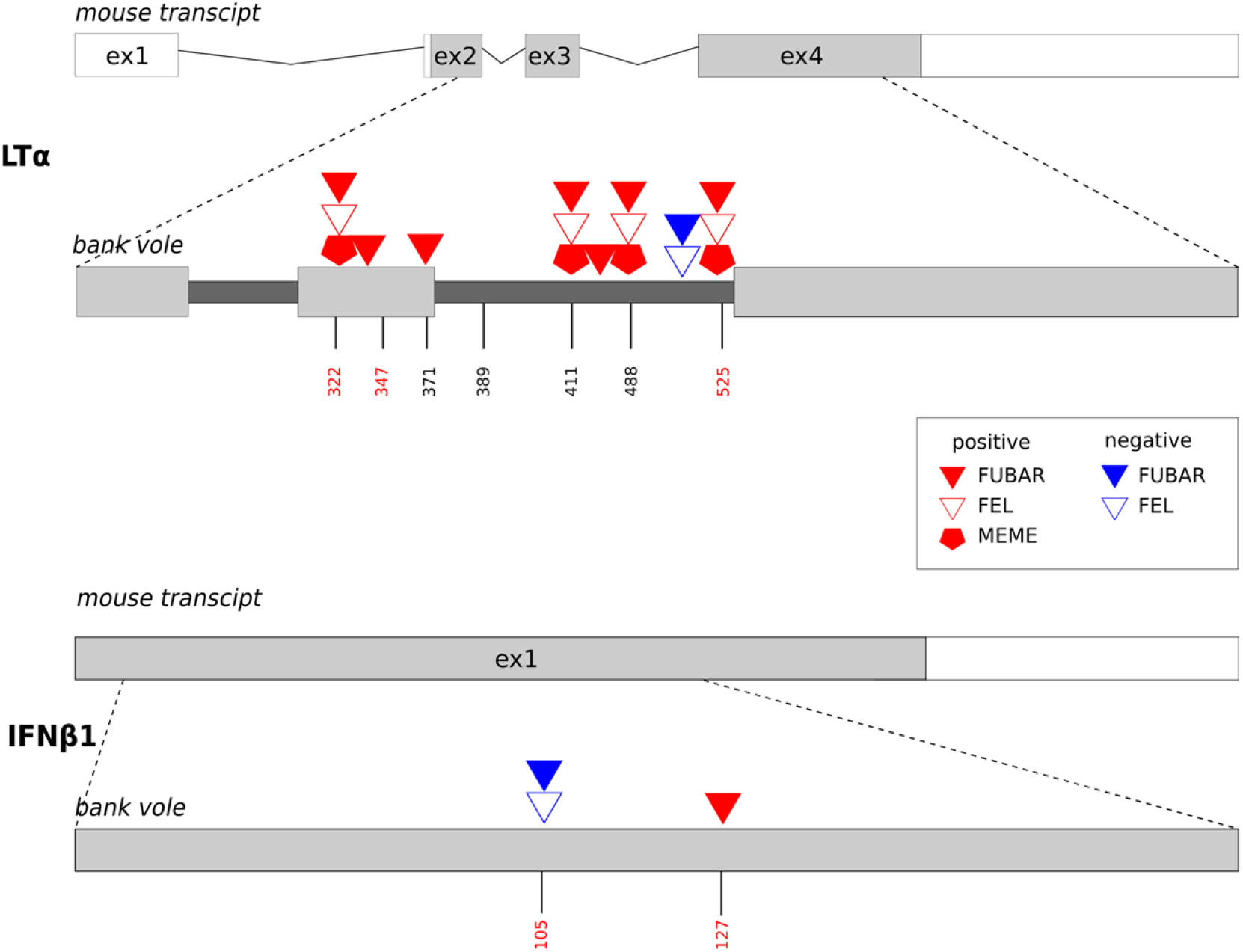
Schematic position of the sites under selection (marked as triangles and pentagons) in LTα and IFNβ1 in relation to exon-intron structure. Positions of SNPs are given with bars, and SNPs significantly associated with parasite load are marked with red.

## 4. Discussion

Studies of associations between polymorphisms in the immunity genes and susceptibility to diseases in wild mammals usually focus on receptor proteins, such as MHC or TLR. Since they physically interact with pathogen-derived motifs, their nucleotide composition can be directly attributed to functional variation. In the case of cytokines - excreted molecules involved in signal transduction – identifying potential targets for selection is not straightforward. Some authors suggested that molecules of such a function are primary affected by purifying selection (Chapman et al. 2016), but several studies in free-living mammals showed otherwise (Turner et al. 2011, 2012). Numerous human studies associated SNP variation in cytokines with a variety of diseases and conditions (reviewed in Hollegaard and Bidwell 2006), and in the present study we show that cytokine variance play also a role in susceptibility to infections in free-living rodents.

We found signatures of positive selection in two variants: LTα 525 and IFNβ1 127 what suggests evolutionary pressure favouring these sites. This may be explained by significant effect of these variants on parasite load what is consistent with a pathogen-mediated model of evolution: repeated rounds of positive selection interspersed with purifying selection. Interestingly, variant IFNβ1 105 associated with susceptibility to two pathogens displayed signatures of purifying selection. Individuals with the genotype TT were more susceptible to the nematode *H. glareoli* but less to blood parasite *Bartonella sp*. compared to other genotypes, and the frequency of the TT genotype was twice lower than the alternative CC. We hypothesize that TT genotype is negatively selected what may be caused by a stronger fitness effect of an infection with bacteria that attacks endothelium and erythrocytes compared to nematodes dwelling the intestine lumen.

Variant IFNβ1 105 affecting parasite load is synonymous. This result may seem confusing, as synonymous substitutions do not affect the amino-acid composition of a protein. However, site-specific signals of selection in this locus strongly suggest that non-neutral pressure is exerted on these sites. The role of non-coding variants may be more significant than previously thought, for instance the list of human diseases associated with synonymous mutations is expanding (Sauna and Kimchi-Sarfaty 2011). In a large meta-analysis of human GWAS, Chen et al. (2010) reported that synonymous SNPs were as often involved in disease mechanisms as non-synonymous SNPs, and they were not in linkage disequilibrium with causal non-synonymous SNPs. Several mechanisms may explain the role of synonymous mutations. They may affect mRNA splicing and stability of transcripts (Cartegni et al. 2002). Although coding for the same amino-acid, some variants may be preferred during the elongation, promoting co-evolution to optimise translation efficiency (Plotkin and Kudla 2011). On the other hand, since synonymous mutations are assumed to be evolutionary silent, their effects might have been under-reported (Chen et al. 2010, Sauna and Kimchi-Sarfaty 2011). This further underlines the importance of studies on synonymous SNPs in non-model species for better understanding processes maintaining genetic diversity within the immune system in the wild.

Another type of non-coding SNPs that we found to significantly affect the parasite load was an intronic variant LTα 525. Notably, it was not linked to any exonic variant, and strong LD was found only to another intronic SNP. Again, human association studies confirmed the role of intronic polymorphisms in susceptibility to diseases (Manolio et al. 2009; Cooper 2010), particularly when mutations are located close to intron-exon junctions or within a branchpoint sequence (Královicová et al. 2006). Intronic variants may also affect expression through alternative splicing or interactions with regulatory elements (Cooper 2010). The intronic LTαvariant 525 that affected susceptibility to infection with *H. mixtum* was located only 18bp from intron-exon junction. Introns in LTα gene were shown to affect expression in several in vitro and in vivo studies (reviewed in Yokley et al. 2010), and intronic SNPs can still affect splicing or expression, even if separated by over 30bp from any splice site (Coulombe-Huntington et al. 2009; Cooper 2010).

Studies of the effect of intronic variants on resistance against infections in the wild are rare but in humans, positively selected SNP associated with susceptibility to Lassa virus were found in interleukin IL21 gene outside the open-reading frame (Andersen et al. 2012). The authors suggested that those variants may lead to regulatory changes such as differential gene expression. Intronic variants may also affect cytokine interactions with other components with the immune system. For instance, an intronic variant in the human IFNγ gene coincides with a putative NF-κ B binding site which might have functional consequences for the transcription of the human IFNγ gene (Pravica et al. 2000). In bank voles, SNP located within the promoter of the TNF gene affected susceptibility to Puumala virus (PUUV) (Guivier et al. 2010). Unfortunately, we could not confirm this pattern as the fragment amplified in the current study did not span 5’ upstream region.

Our finding suggests that intronic and synonymous variants may play an important role in susceptibility to infections in wild systems. Thus, it should not be neglected in studies of the genetic background of host-parasite co-evolution.

## Author contributions

AK conceived the study, genotyped samples and analysed data, ABi did test of selection. AK and ABi prepared the manuscript. AK and ABa identified helminth infections and collected data in the field with the help from EM. EM and RWF analysed infections with blood parasites.

## Acknowledgments

We thank D.R. Laetsch and M.A. Wenzel for their valuable hints on the bioinformatic pipeline and data analysis. We are thankful to W. Babik who provided access to an Illumina MiSeq platform, and to K. Dudek who prepared the Nextera library.

## References

Andersen KG, Shylakhter I, Tabrizi S, Grossman SR, Happi CT, Sabeti PC (2012). Genome-wide scans provide evidence for positive selection of genes implicated in Lassa fever. Philos Trans R Soc Lond B Biol Sci., 367: 868–877

Babik W, Dudek K, Fijarczyk A, Pabijan M, Stuglik M, Szkotak R, Zieliński P (2015). Constraint and Adaptation in newt Toll-Like Receptor Genes. Genome Biolo Evol, 7: 81–95.

Bajer A, Behnke JM, Pawełczyk A, Kuliś K, Sereda MJ, Siński E (2005) Medium-term temporal stability of the helminth component community structure in bank voles (*Clethrionomys glareolus*) from the Mazury Lake District region of Poland. Parasitology 130: 213–228.

Barbier M, Delahaye NF, Fumoux F, Rihet P. (2008) Family-based association of a low producing lymphotoxin-alpha allele with reduced Plasmodium falciparum parasitemia. Microbes Infect 10: 673–679.

Benjamini Y, Yekutieli D. (2005) False discovery rate–adjusted multiple confidence intervals for selected parameters. J Am Stat Assoc, 100: 71–81.

Cartegni L, Chew SL, Krainer, AR. (2002). Listening to silence and understanding nonsense: exonic mutations that affect splicing. Nature Rev Genet 3: 285–298

Castelli EC, Mendes-Junior CT, Sabbagh A, Porto IO, Garcia A, Ramalho J, Lima TH, Massaro JD, Dias FC, Collares CV, Jamonneau V, Bucheton B, Camara M, Donadi E (2015). HLA-E coding and 3’ untranslated region variability determined by next-generation sequencing in two West-African population samples. Hum Immunol. 76: 945–53.

Chapman JR, Hellgren O, Helin AS, Kraus RH, Cromie RL, Waldenström J. (2016). The Evolution of Innate Immune Genes: Purifying and Balancing Selection on β-Defensins in Waterfowl. Mol Biol Evol, 33: 3075–3087.

Chen R, Davydov EV, Sirota M, Butte AJ (2010) Non-Synonymous and Synonymous Coding SNPs Show Similar Likelihood and Effect Size of Human Disease Association. PLoS ONE 5: e13574.

Cooper DN. (2010). Functional intronic polymorphisms: Buried treasure awaiting discovery within our genes. Hum Genomics 4: 284–288.

Coulombe-Huntington J, Lam KCL, Dias C, Majewski J. (2009). Fine-scale variation and genetic determinants of alternative splicing across individuals. PLoS Genet 5:e1000766.

De Togni P, Goellner J, Ruddle NH, Streeter PR, Fick A, Mariathasan S, Smith SC, Carlson R, Shornick LP, Strauss-Schoenberger J. (1994). Abnormal development of peripheral lymphoid organs in mice deficient in lymphotoxin. Science 264:703–707.

Delport W, Poon AFY, Frost SDW, Kosakovsky Pond SL. (2010). Datamonkey 2010, a suite of phylogenetic analysis tools for evolutionary biology. Bioinformatics, 26: 2455–2457.

Ferrer-Admetlla A, Bosch E, Sikora M, Marquès-Bonet T, Ramírez-Soriano A, Muntasell A, Navarro A, Lazarus R, Calafell F, Bertranpetit J, Casals F. (2008). Balancing Selection Is the Main Force Shaping the Evolution of Innate Immunity Genes. J Immunol, 181: 1315–1322.

Fornůsková A, Vinkler M, Pagès M, Galan M, Jousselin E, Cerqueira F, Morand S, Charbonnel N, Bryja J, Cosson JF (2013). Contrasted evolutionary histories of two Toll-like receptors Tlr4 and Tlr7 in wild rodents Murinae. BMC Evol Biol, 13: 194.

Garrison E, Marth G. (2012). Haplotype-based variant detection from short-read sequencing. arXiv preprint arXiv:1207.3907.

Guivier E, Galan M, Salvador AR, Xuéreb A, Chaval Y, Olsson GE, Essbauer S, Henttonen H, Voutilainen L, Cosson JF, Charbonnel N. (2010). Tnf-α expression and promoter sequences reflect the balance of tolerance/resistance to Puumala hantavirus infection in European bank vole populations. Infect Genet Evol 10: 1208–1217.

Henricksen S, Pohlenz J (1981) Staining of cryptosporidia by modified Ziehl–Nielsen technique. Acta Vet Scand 22: 594–596.

Hollegaard MV, Bidwell JW. (2006). Cytokine gene polymorphism in human disease: on-line databases Genes Immun 7: 269–276

Hong EP, Park JW. (2012). Sample size and statistical power calculation in genetic association studies. Genomics Inform 10: 117–22.

Iizuka K, Chaplin DD, Wang Y, Wu Q, Pegg LE, Yokoyama WM, Fu YX. (1999). Requirement for membrane lymphotoxin in natural killer cell development. Proc Natl Acad Sci USA 96: 6336–6340.

Jepson A, Banya W, Sisay-Joof F, Hassan-King M, Nunes C, Bennett S, Whittle H. (1997). Quantification of the relative contribution of major histocompatibility complex (MHC) and non-MHC genes to human immune responses to foreign antigens. Infect Immun 65: 872–876.

Kloch A, Bierzycka A. (2020). Post-glacial phylogeography and variation in innate immunity loci in a sylvatic rodent, bank vole *Myodes glareolus*. Mamm Biol, 100: 141–154

Kloch A, Babik W, Bajer A, Siński E, Radwan J. (2010). Effects of an MHC-DRB genotype and allele number on the load of gut parasites in the bank vole Myodes glareolus. Mol Ecol 19 Suppl 1:255–265.

Kloch A, Wenzel MA, Laetsch DR, Michalski O, Bajer A, Behnke J, Welc-Falęciak R, Piertney, SB (2018). Signatures of balancing selection in toll-like receptor (TLR) genes – novel insights from a free-living rodent. Sci Rep 8: 8361.

Kosakovsky Pond SL, Frost SD. (2005). Not so different after all, a comparison of methods for detecting amino acid sites under selection. Mol Biol Evol 22: 1208–1222.

Kosakovsky Pond SL, Posada D, Gravenor MB, Woelk CH, Frost SDW (2006). GARD: a genetic algorithm for recombination detection. Bioinformatics, 22: 3096–3098

Královicová J, Lei H, Vorechovský I. (2006). Phenotypic consequences of branch point substitutions. Hum Mutat 27: 803–813.

Li H. (2013). Aligning sequence reads, clone sequences and assembly contigs with BWA-MEM. arXiv preprint arXiv:1303.3997.

Lima THA, Buttura RV, Donadi EA, Veiga-Castelli LC, Mendes-Junior CT, Castelli E. (2016). HLA-F coding and regulatory segments variability determined by massively parallel sequencing procedures in a Brazilian population sample. Hum Immunol. 76: 841–853.

Manolio TA, Collins FS, Cox NJ, Goldstein DB, Hindorff LA, Hunter DJ, McCarthy MI, Ramos EM, Cardon LR, Chakravarti A, et al. (2009). Finding the missing heritability of complex diseases. Nature 461: 747–753.

Murphy K, Travers P, Walport M, Janeway, C. (2012). Janeway’s immunobiology. 8th ed. New York: Garland Science.

Murrell B, Moola S, Mabona A, Weighill T, Sheward D, Kosakovsky Pond SL, Scheffler K (2013). FUBAR, a fast, unconstrained bayesian approximation for inferring selection. Mol Biol Evol, 30: 1196–1205.

Murrell B, Wertheim JO, Moola S, Weighill T, Scheffler K, Kosakovsky Pond SL (2012). Detecting individual sites subject to episodic diversifying selection. PLoS Genetics, 8: e1002764.

Nagarajan UM (2011) Induction and Function of IFNβ During Viral and Bacterial Infection. Crit Rev Immunol. 31: 459–474.

Nielsen R, Paul JS, Albrechtsen A, Song YS. (2011). Genotype and SNP calling from nextgeneration sequencing data. Nat Rev Genet 12:443–451.

Norman AF, Regnery R, Jameson P, Greene C, Krause DC (1995). Differentiation of Bartonella-like isolates at the species level by PCR-restriction fragment length polymorphism in the citrate synthase gene. J Clin Microbiol 33: 1797–1803.

Persing DH, Mathiesen D, Marshall WF, Telford SR, Spielman A, Thomford JW, Conrad PA (1992). Detection of *Babesia microti* by polymerase chain reaction. J Clin Microbiol, 30: 2097–2103.

Plotkin JB, Kudla G. (2011) Synonymous but not the same: the causes and consequences of codon bias. Nature Rev Genet 12: 32–42

Pravica V, Perrey C, Stevens A, Lee JH, Hutchinson IV. (2000) A single nucleotide polymorphism in the first intron of the human IFN-gamma gene: absolute correlation with a polymorphic CA microsatellite marker of high IFN-gamma production. Hum Immunol 61: 863–6

Purcell S, Neale B, Todd-Brown K, Thomas L, Ferreira MAR, Bender D, Maller J, Sklar P, de Bakker PIW, Daly MJ & Sham PC (2007) PLINK: a toolset for whole-genome association and population-based linkage analysis. Am J Hum Genet 81: 559–575.

R Core Team (2020). R: A language and environment for statistical computing. R Foundation for Statistical Computing, Vienna, Austria. https://www.R-project.org/.

Radwan J, Babik W, Kaufman J, Lenz TL, Winternitz J. 2020. Advances in the Evolutionary Understanding of MHC Polymorphism. Trends Genet, 36: 298–311

Sauna ZE, Kimchi-Sarfaty Ch. (2011). Understanding the contribution of synonymous mutations to human disease. Nat Rev Genet 12: 683–691

Turner AK, Begon M, Jackson JA, Bradley JE, Paterson S (2011) Genetic Diversity in Cytokines Associated with Immune Variation and Resistance to Multiple Pathogens in a Natural Rodent Population. PLoS Genet 7: e1002343.

Turner AK, Begon M, Jackson JA, Paterson S. (2012). Evidence for selection at cytokine loci in a natural population of field voles (*Microtus agrestis*). Mol Ecol 21:1632–1646.

Ware CF. (2005). Network communications: lymphotoxins, LIGHT, and TNF. Annu Rev Immunol 23: 787–819.

Welc-Faleciak R, Paziewska A, Bajer A, Behnke JM, Siński E. (2008). Bartonella spp. infection in rodents from different habitats in the Mazury Lake District, Northeast Poland. Vector Borne Zoonotic Dis 8:467–474.

Yokley BH. (2010). Regulation of the lymphotoxin alpha gene: characterization of elements located between the transcription and translation start sites that impact expression. PhD thesis, Georgetown University

